# Fusion of Complement Fragment C3d Enhances Germinal Center Responses to HIV-1 Envelope Glycoproteins

**DOI:** 10.1101/2025.06.28.661730

**Authors:** Marlon de Gast, Sara Hernández-Pérez, Arpan Pradhan, Sabine Kruijer, Wouter Olijhoek, Isabel J. L. Baken, Christoph Matti, Martijn Breeuwsma, Zhang Sung Tean, Willemijn J. C. van Keizerswaard, Samantha Zoomer, Judith A. Burger, Tom G. Caniels, Kwinten Sliepen, Ronald Derking, Manoj Thapa, Arash Grakoui, Balthasar A. Heesters, Mathieu Claireaux, Rogier W. Sanders, Pavel Tolar, Sudhir P. Kasturi, Marit J. van Gils

**Author notes:** Authors contributed equally (Sudhir P. Kasturi, Marit J. van Gils). Correspondence to &.

## Abstract

Eliciting sustained germinal center (GC) responses is critical for the development of an effective HIV-1 vaccine, yet HIV-1 envelope glycoprotein (Env) immunogens often fail to elicit GC responses required for the maturation of cognate B cells that secrete broadly neutralizing antibodies (bNAbs). Effective antigen recognition is important for initial B cell priming, activation, and GC engagement. Since complement opsonization contributes to antigen recognition, we investigated whether C3d fusion could enhance the GC response of the stabilized HIV-1 Env immunogen based on a consensus sequence (ConM Env). Our results demonstrate that ConM Env-C3d induced potent HIV-specific B cell activation in vitro compared to ConM Env alone. We also observed that the C3d fusion enhanced antigen presentation by human tonsil-derived follicular dendritic cells (FDCs) to HIV-specific B cell lines. Moreover, mouse immunization studies combining ConM Env-C3d with the AddaS03 adjuvant revealed significantly enhanced early GC formation and prolonged antigen display and retention on FDCs for up to 56 days, highlighting improved antigen persistence within GCs. These immunological enhancements, including a more focused early antibody response, correlated with improved virus neutralization. Additionally, we observed sex-based differences in immune responses, with female mice showing stronger antibody responses and enhanced antigen retention compared to males. These findings suggest that C3d fusion can enhance GC engagement and improve the immunogenicity of HIV-1 vaccines.

## Introduction

Developing a broad and effective vaccine against HIV-1 remains a persistent challenge in vaccinology. The HIV-1 envelope glycoprotein (Env) is the only target for neutralizing antibodies (NAbs) but presents significant challenges as a vaccine immunogen due to its extreme genetic variability and dense glycan shield. While stabilization of the Env trimer, for example through SOSIP-based design *(Sanders et al. 2013, 2015; Sanders and Moore 2024)*, and germline B cell targeting strategies *(Caniels et al. 2023, 2025)* have improved Env immunogenicity in both animal models and humans, the resulting responses often lack the level of B cell maturation needed to achieve broad and durable protective immunity.

The potency of vaccine-induced antibody responses depends on robust germinal center (GC) reactions, which are specialized microenvironments within secondary lymphoid tissues where B cells undergo affinity maturation *(Lee et al. 2022; Mesin, Ersching, and Victora 2016)*. Within GCs, B cells compete for antigen and mutate their B cell receptors (BCRs). High-affinity B cells are selected with help from follicular dendritic cells (FDCs), which retain intact antigen via immune complexes (ICs) for prolonged periods, and CD4^+^ T follicular helper (Tfh) cells, which provide essential survival signals (*De Silva and Klein 2015; Aung et al. 2023*). Through the introduction of mutations, this iterative selection process not only improves affinity but also promotes antibody breadth by favoring expansion and survival of B cells capable of recognizing variant forms of the antigen *(Abbott et al. 2018)*. Sustained GC activity is particularly important in this context, as prolonged B cell evolution is often required to generate the breadth and potency needed to neutralize diverse HIV-1 strains *(Victora and Nussenzweig 2022; Lee et al. 2022)*.

Despite advances in Env design, HIV-1 immunogens often fail to induce GC responses of sufficient magnitude or duration to support the development of broad neutralizing antibodies (bNAbs). This limitation stems from Env’s poor immunogenicity, including its limited ability to engage B cells effectively and drive sustained GC activity *(Martin et al. 2020; Abbott et al. 2018)*. Although recent studies in non-human primates (NHPs) have reported improvements in antibody durability and potency, the responses remain suboptimal *(Cirelli et al. 2019; Martin et al. 2020; Lee et al. 2022)*. Achieving long-term vaccine efficacy requires enhanced GC responses to support the maturation towards durable bNAbs. These findings highlight the need for strategies that enhance GC engagement and maintenance and support bNAb maturation in response to Env-based vaccines.

To overcome the limited GC engagement by HIV-1 Env immunogens, strategies that enhance early B cell activation and improve antigen retention within GCs are needed. One promising strategy is to leverage the immune system’s own co-stimulatory molecules, such as the complement fragment C3d, which has adjuvant-like properties. C3d is produced during complement activation, following cleavage of factor C3 *(Lachmann, Pangburn, and Oldroyd, 1982)*. When the complement cascade is triggered by foreign antigens, C3d is deposited onto their surface, marking them for recognition by complement receptor 2 (CR2, also known as CD21), which is expressed on both B cells and follicular dendritic cells (FDCs) *(Szakonyi et al. 2001; Dempsey et al. 1996)*. This interaction plays a dual role: it enhances long-term antigen display by FDCs and simultaneously acts as a co-stimulatory signal for B cells, lowering their activation threshold and boosting responsiveness. When covalently fused to antigens, C3d amplifies B cell activation by simultaneously engaging the B cell receptor and CR2, a mechanism shown to enhance activation by up to 10,000-fold in model systems *(Dempsey et al. 1996)*. This property is especially valuable for weak immunogens like HIV-1 Env, which often struggle to activate low-affinity B cells. Previous studies have demonstrated that C3d fusion increases antibody titers and improves affinity maturation and viral neutralization against HIV-1 Env proteins, including gp120 and native flexible linked (NFL) trimers *(Bale et al. 2023; Bower and Ross 2006)*. However, the specific impact of C3d fusion on GC dynamics remains poorly understood.

In addition to strong initial activation, robust GC reactions rely on sustained antigen availability. B cells must repeatedly acquire intact antigen, process it, and present its peptides on MHC II molecules to receive continued help from Tfh cells and compete effectively in the light zone *(Aung et al. 2023; Martínez-Riaño et al. 2023; Lee et al. 2022)*. Yet, recombinant HIV-1 Env trimers do not readily accumulate in GC follicles following conventional immunization *(Martin et al. 2020)*. Strategies such as slow-delivery immunization have partially addressed this by improving GC responses through sustained antigen availability and reduced targeting of immunodominant, non-neutralizing epitopes *(Cirelli and Crotty 2017; Cirelli et al. 2019)*. Given that FDCs retain complement-opsonized ICs via CR2 and Fcγ receptors, fusing C3d to Env may enhance its display and retention within GCs *(Aung et al. 2023; Heesters, Myers, and Carroll 2014; Heesters et al. 2021)*. However, the effect of C3d fusion on the retention of the antigen within the GC has not yet been investigated. By promoting both B cell activation and extended antigen availability, C3d fusion may represent a promising strategy to overcome persistent challenges in Env-based vaccine design.

In this study, we aimed to enhance the GC response of Env glycoproteins through C3d fusion. We used a native-like Env trimer immunogen, ConM SOSIP.v7 Env (ConM Env), which is based on an artificial consensus sequence of all HIV-1 isolates in group M. This immunogen has consistently induced robust autologous NAb responses in NHPs *(Sliepen et al. 2019)* and, more recently, in humans *(Reiss et al., under revision; Pollock et al. 2025). In vitro*, we observed increased B cell activation and enhanced antigen presentation by human FDCs to HIV-specific B cells when ConM Env was fused with C3d, underscoring the direct effects of C3d on B cell and FDC function.*In vivo*, C3d-conjugated ConM Env elicited higher binding antibody titers, increased virus neutralization titers, and enhanced GC B cell responses in mice when administered with the adjuvant AddaS03. Importantly, we observed enhanced antigen retention in draining lymph nodes after immunization, suggesting that C3d fusion promotes prolonged antigen availability in GCs, further supporting its role in facilitating robust GC responses. Interestingly, we also observed sex-based differences, with female mice showing stronger antibody responses and antigen retention compared to males, suggesting that gender may influence the effectiveness of C3d fusion in modulating immune responses. Overall, this study demonstrates that C3d fusion is a promising strategy for optimizing vaccine responses by enhancing B cell engagement and sustaining GC activity of HIV-1 Env-based immunogens.

## Results

### C3d Fusion to HIV-1 ConM Env Immunogen: Structural Integrity and Base Epitope Masking

To enhance the GC response to HIV-1 Env glycoproteins, we investigated the impact of fusing the complement fragment C3d to ConM Env, a native-like immunogen derived from a consensus sequence of HIV-1 group M *(Sliepen et al. 2019)*. We generated two ConM Env-C3d constructs: one incorporating human C3d and one incorporating murine C3d, each genetically fused to the C-terminus of the gp140 subunits using a twelve–amino acid GSG-linker (Fig. 1A). These constructs were designed to function optimally in both human and mouse cell systems (Fig. S1).

**Figure 1:**
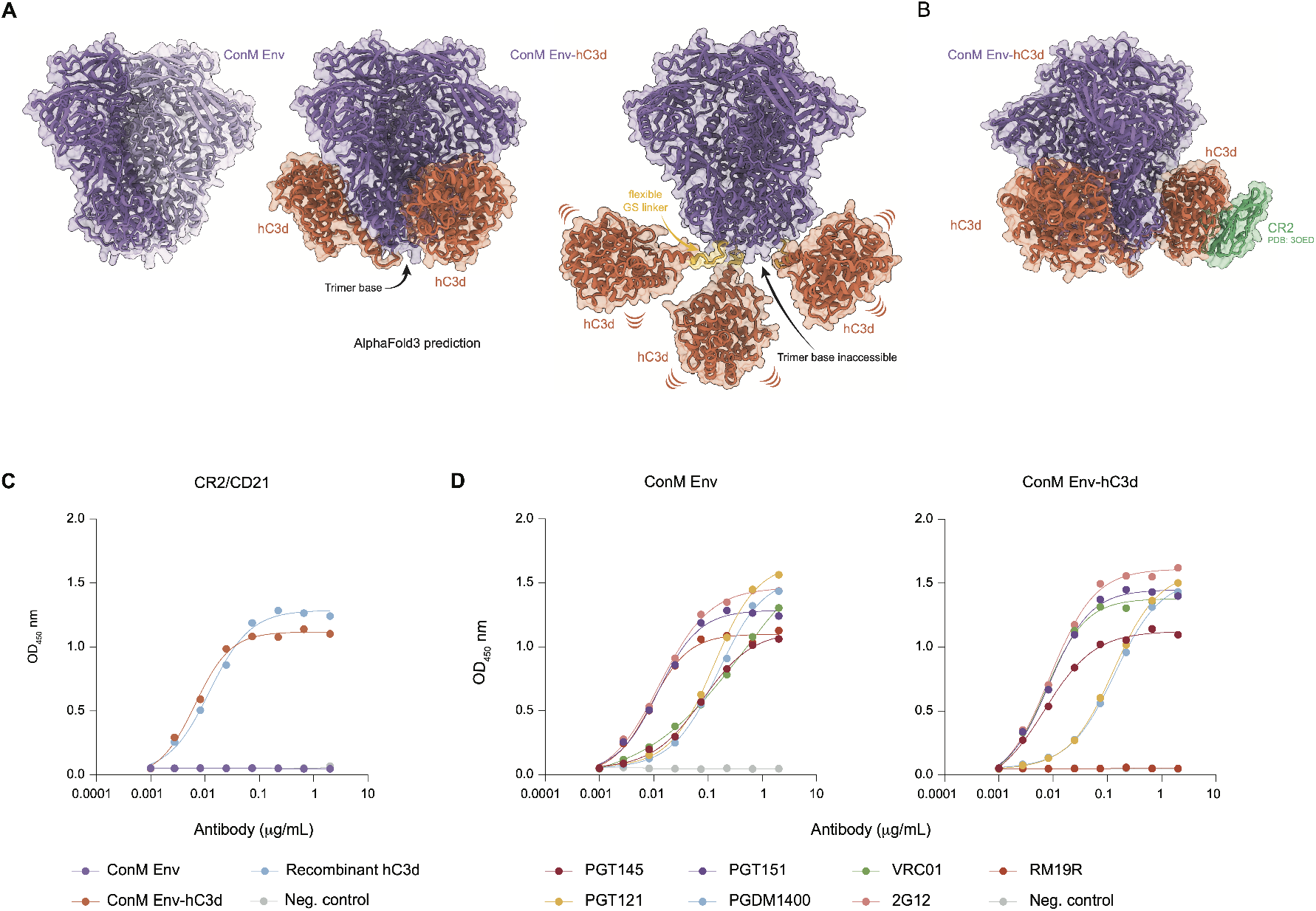
Conformation and antibody binding of ConM Env and ConM Env-hC3d. **(A)** AlphaFold3 prediction of the ConM Env (purple)-hC3d (orange) fusion construct. The trimer base epitope is indicated with an arrow. The flexible glycine-serine (GS) linker (yellow) may allow for rotation of the hC3d subunits, potentially obscuring immunodominant trimer base epitopes. **(B)** Superposition of ConM Env-hC3d onto a hC3D-complement receptor 2 (CR2) structure (PDB: 3OED). The predicted angle of the hC3d domain allows for CR2 binding. **(C)** ELISA binding assay of complement receptor 2 (CR2)/CD21 with ConM Env, ConM Env-hC3d, recombinant hC3d, and the SARS-CoV-2 Wuhan Spike as a negative control. **(D)** ELISA binding curves for antibodies PGT145, PGT121, PGT151, PGDM1400, VRC01, 2G12, RM19R against the ConM Env (left) and ConM Env-hC3d (right) constructs, with SARS-CoV-2-binding mAb COVA1-16 as negative control.

To evaluate the predicted geometry of the fusion, we used AlphaFold3 to model the ConM Env-hC3d trimer. The prediction suggests that the three C3d domains cluster near the trimer base, where their flexible linkers may permit rotation that partially shields the base epitopes while preserving CR2 accessibility.

We confirmed robust binding of CR2 to ConM Env-hC3d (Fig. 1B), demonstrating proper folding of the C3d moiety and its potential to engage with the complement receptor (Fig. 1C). Additionally, our results showed that bNAbs, such as PGDM1400 (V2-apex), VRC01 (CD4-binding site), PGT121 and 2G12 (high mannose patch-directed), and PGT145, and PGT151 (quaternary-dependent bNAbs), bound to both ConM Env and ConM Env-hC3d (Fig. 1D). These findings demonstrate that C3d fusion does not affect trimer folding or epitope accessibility.

Remarkably, the non-neutralizing antibody RM19R bound to ConM Env but not to ConM Env-hC3d (Fig. 1D), suggesting decreased accessibility of the trimer base, possibly due to steric hindrance by the fused C3d domains. Thus, C3d fusion may reduce the immunodominance typically associated with Env-induced base-specific antibody responses.

### ConM Env-C3d Enhances Activation of Human HIV-1 Env-specific B Cells *in Vitro*

To assess the impact of C3d fusion on BCR-mediated B cell activation, we utilized HIV-1 Env-specific human Ramos B cells engineered to stably express the bNAbs PGDM1400 or PGT121 as their BCRs and measured their activation through Ca^2+^ influx *in vitro*. We evaluated three protein constructs: ConM Env, ConM Env-hC3d (the human C3d-fused version), and ConM Env supplemented with soluble hC3d, all tested at various equimolar concentrations.

PGT121-expressing B cells showed robust activation in response to ConM Env-hC3d across all tested concentrations (10, 1, 0.1, 0.01, and 0.001 µg/ml Env-equivalent; Fig. 2). In contrast, ConM Env alone triggered markedly weaker activation at lower concentrations and showed no measurable activation above baseline at 0.01 µg/ml. At this concentration, ConM Env-hC3d elicited an ~90-fold stronger response than ConM Env alone (AUC 442 vs. 5; Fig. S2A). ConM Env supplemented with soluble hC3d resulted in intermediate activation levels, falling between those of ConM Env alone and ConM Env-hC3d, but also failed to induce responses at the lowest doses (Fig. 2). These findings highlight the enhanced efficacy of C3d fusion to activate B cells compared to admixed supplementation with soluble C3d.

**Figure 2:**
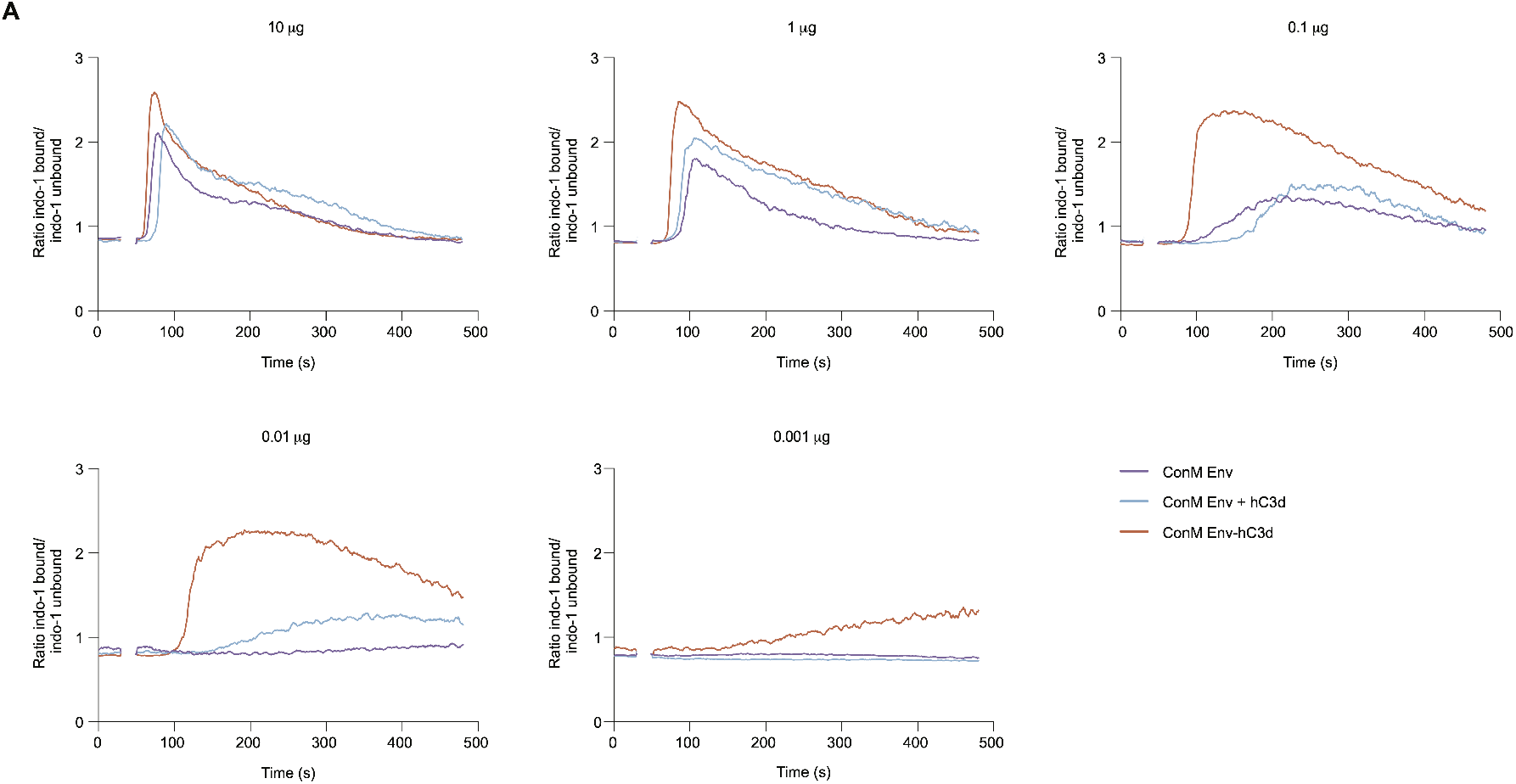
Activation of B cells expressing bNAb PGT121 by ConM Env and ConM Env-hC3d. Ramos B cell activation of PGT121 IgG B cells as measured by calcium (Ca^2+^) flux assay using equimolar amounts of ConM Env, ConM Env-hC3d, or ConM Env supplemented with recombinant hC3d (ConM Env + hC3d) at concentrations of 10 µg, 1 µg, 0.1 µg, 0.01 µg, and 0.001 µg (Env-equivalent mass). ConM Env and recombinant hC3d were co-administered at a molar ratio matching the stoichiometry of ConM Env-hC3d. A baseline without antigen was established between 0–30 s, after which antigen was added to the B cells (30–50 s).

Similarly, PGDM1400-expressing B cells displayed distinct activation profiles in response to the different constructs. While ConM Env alone failed to activate these cells (Fig. S2C), both ConM Env-hC3d and ConM Env supplemented with soluble hC3d induced activation at 10 µg. Among the conditions, ConM Env-hC3d elicited the strongest response. At 1 µg, only the fused construct continued to induce detectable activation, whereas the response from soluble hC3d was completely lost. At lower concentrations (≤ 0.1 *µ*g), none of the constructs induced measurable activation. Control experiments using ionomycin (positive control) and SARS-CoV-2 spike (non-cognate antigen) confirmed the specificity of the assay (Figs. S2B and S2C).

These results demonstrate that the covalent linkage of C3d to ConM Env significantly enhances B cell sensitivity to BCR-mediated activation, supporting more B cell activation at lower antigen concentrations. This effect may extend to lower-affinity B cell receptors *in vivo*, where signaling thresholds are typically higher.

### Enhanced Presentation of ConM Env-hC3d by Human FDCs to Antigen-specific B Cells

Given the role of FDCs in antigen retention within germinal centers, we hypothesized that fusing C3d to the ConM Env immunogen would enhance its capture by FDCs. Because FDCs capture immune complexes via CR2 and Fcγ receptors, we compared binding of fluorescently labeled ConM Env and ConM Env-hC3d to FDCs, testing direct antigen binding as well as immune complexes (ICs; ConM Env-IC, ConM Env-hC3d-IC) and complement-opsonized immune complexes (CO-ICs; ConM Env CO-IC, ConM Env-hC3d CO-IC). Human FDCs were isolated from palatine tonsils, cultured, and identified as PDPN^+^, CD55^+^, CD21^+^, and CD35^+^cells. As expected, only a small percentage of FDCs captured the ConM Env protein alone (7.0%; Fig. 3A). Binding increased when delivered as immune complexes, both with and without complement opsonization: ConM Env-IC (33%) and ConM Env CO-IC (48%). In contrast, ConM Env-hC3d showed significantly higher capture FDCs, both in free form (69%; p = 0.003), and when presented as ConM Env-hC3d-IC (85%; p = 0.025) or ConM Env-hC3d CO-IC (94%; p = 0.0009; Fig. 3A).

**Figure 3:**
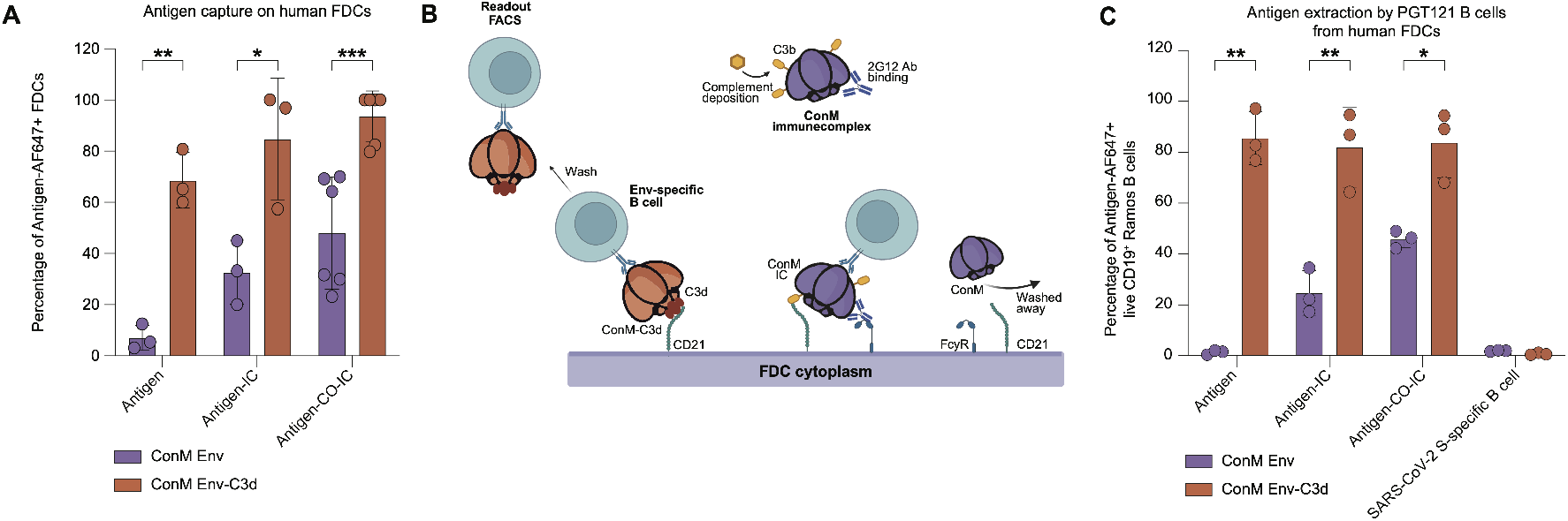
FDC-mediated presentation of ConM Env-C3d to HIV-1 Env-specific B cells. **(A)** Percentage of live PDPN^+^ CD21^+^ CD35^+^ FDCs positive for binding ConM Env-AF647 or ConM Env-hC3d-AF647, presented either as the antigen alone, within an immune complex (IC; 2G12 mAb), or within a complement-opsonized immune complex (CO-IC; active human serum + 2G12 mAb). **(B)** Schematic of antigen extraction by PGT121 B cells from human FDCs. Fluorescently labeled antigen, either alone or within an immune complex (IC), binds to FDCs via C3d/C3b–CD21 interactions or antibody Fc–FcγR engagement. After incubation, unbound antigen is removed by washing. HIV-specific PGT121 Ramos B cells are then added, followed by a second wash. Antigen extraction is assessed by flow cytometry, measuring fluorescent signal on HIV-1 Env-specific B cells. Schematic was created with BioRender.com. **(C)** Percentage of live CD19^+^ GFP^+^ PGT121-expressing Ramos B cells positive for binding ConM Env-AF647 or ConM Env-hC3d-AF647, presented by FDCs as described in the schematic of Figure 5B. Antigen was presented either alone, within an immune complex (IC; 2G12 mAb), or as a complement-opsonized IC (CO-IC; active human serum + 2G12 mAb). A SARS-CoV-2 spike-specific Ramos B cell line (COVA1-16) was included as a negative control to assess antigen specificity in the antigen-CO-IC condition. Bar plots depict the mean ± SD. Each dot represents one donor; paired comparisons were performed using two-tailed paired t-tests. Significance is indicated as: ns, not significant (p > 0.05); *, p < 0.05; **, p < 0.01; ***, p < 0.001.

To further investigate the role of C3d fusion in antigen retention by FDCs and subsequent extraction by B cells, we examined its impact on antigen transfer from FDCs to human HIV-specific Ramos B cells (PGT121-expressing BCR; Fig. 3B). Since only the antigens retained on FDCs can be extracted by B cells, lower binding to FDCs resulted in reduced antigen availability for extraction. In the absence of ICs, only ConM Env-hC3d (87%) was efficiently extracted by PGT121 B cells, while ConM Env remained nearly undetectable (1.0%; p = 0.004; Fig. 3C). IC formation modestly improved antigen capture of ConM Env (25% for ConM Env-IC and 46% for ConM Env CO-IC), but uptake remained significantly lower than ConM Env-hC3d-IC (83%; p = 0.008) and ConM Env-hC3d CO-IC (85%; p = 0.025; Fig. 3C). As a specificity control, we used a SARS-CoV-2 spike-specific (COVA1-16) Ramos B cell line, which showed no capture of any of the antigens and their complexes (Fig. 3C). These results demonstrate that C3d fusion to the ConM Env immunogen enhances antigen capture by FDCs and facilitates more efficient extraction by antigen-specific B cells, supporting its potential to improve immunogen retention and display.

### C3d Fusion Enhances Early Antibody Responses and Neutralization Efficacy

Next, we assessed the immunogenicity of ConM Env-C3d trimers in mice, using murine C3d (mC3d) to ensure compatibility with the murine CR2. Mice were immunized at weeks 0, 4, and 8 with either 10 µg of ConM Env or an equimolar amount of ConM Env-mC3d, administered with or without AddaS03, an oil-in-water emulsion-based adjuvant (Fig. 4A). We also monitored and compared immunological responses between male and female mice to identify any variations attributable to sex. After the initial immunization, detectable Ab responses against the ConM Env trimers were observed only in mice receiving ConM Env adjuvanted with AddaS03, with the addition of C3d significantly enhancing the antibody titers further (Fig. 4B; p < 0.0001). Following the first booster at week 8, antibody titers increased further in AddaS03-adjuvanted groups, with ConM Env-mC3d inducing significantly higher ConM Env-specific antibody responses compared to ConM Env with AddaS03 (Fig. 4B; p < 0.0001). After the second booster, antibody responses against ConM Env trimers were evident in all mice, irrespective of adjuvant usage. However, the additional effect of C3d fusion on ConM Env-specific antibody titers diminishes after three immunizations, resulting in similar serum antibody levels between the ConM Env and ConM Env-mC3d groups when male and female mice were combined (Fig. 4B). Interestingly, among female mice specifically, those vaccinated with ConM Env-C3d demonstrated significantly higher antibody titers against ConM Env than those who received only the ConM Env (p = 0.0027; Fig. 4C).

**Figure 4:**
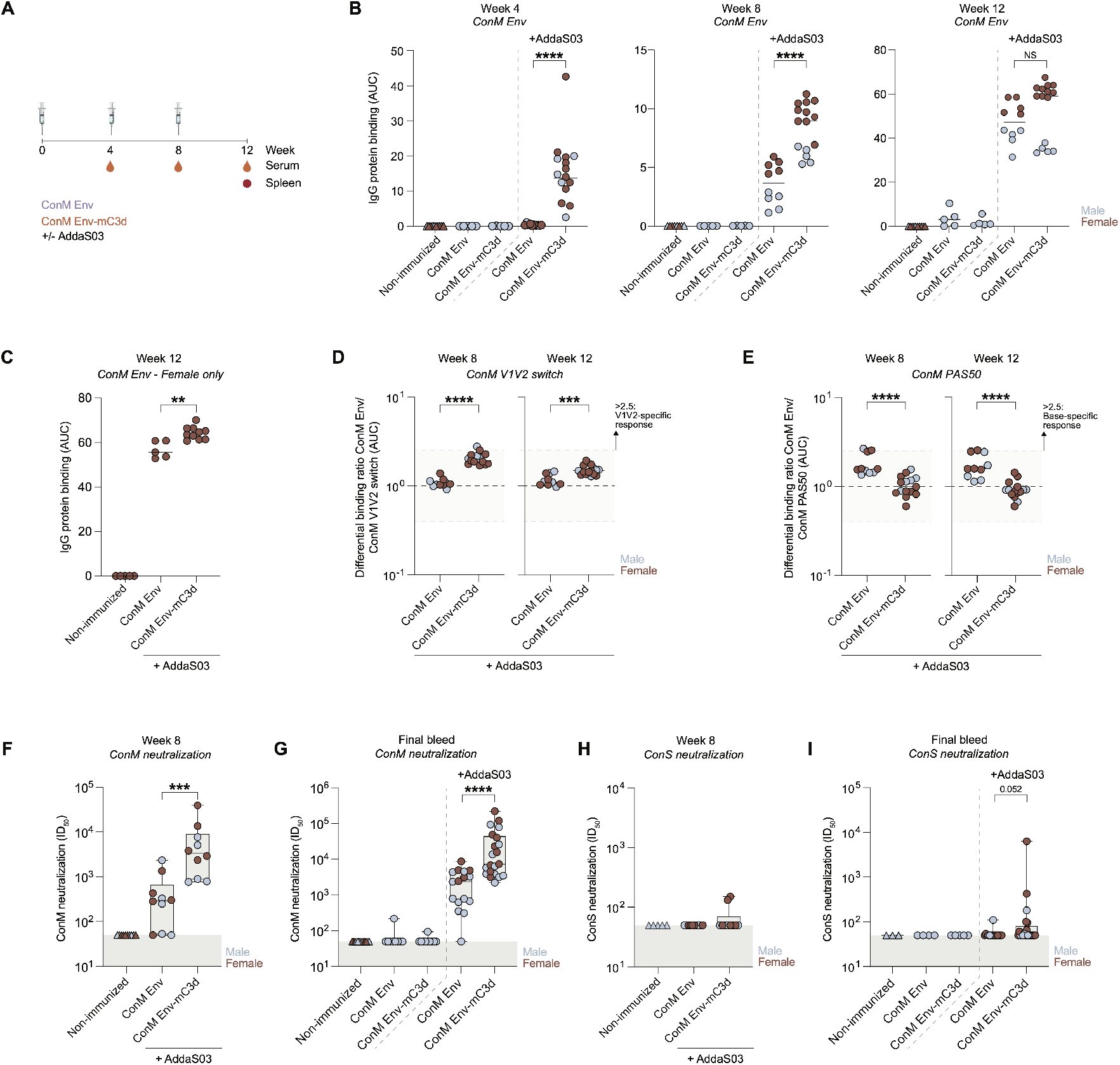
Humoral responses following immunization with ConM Env or ConM Env-mC3d, with or without AddaS03 adjuvant. **(A)** Immunization and sampling schedule. Mice received 10 µg ConM Env or ConM Env-mC3d (equimolar) with or without AddaS03. Serum, lymph nodes (LNs), and spleens were collected at indicated time points. **(B)** ConM Env-specific IgG levels measured by multiplex binding assay at week 4 (left), week 8 (middle), and week 12 (right), plotted as area under the curve (AUC). Data are stratified by sex with male and female mice shown in blue and maroon, respectively. **(C)** ConM Env-specific IgG responses in female mice at week 12, measured by multiplex binding assay and presented as AUC. **(D)** Differential IgG binding ratio of ConM Env to V1V2 switch variant (ConM Env / ConM V1V2 switch), calculated from AUC values at week 8 and 12. Ratios >2.5 (shaded area) indicate a significant V1V2-focused response. **(E)** Differential IgG binding ratio of ConM Env to base-mutant (PAS50) variant (ConM Env / ConM PAS50) at week 8 and 12. Ratios >2.5 (shaded area) indicate a significant base-focused response. (H) Neutralization titers (ID50) against ConS pseudovirus at week 8 and (I) at the final bleed (week 12). Data are stratified by sex, with male (blue) and female (maroon) mice shown separately. Data are shown as scatter plots or box-and-whisker plots (min to max), with medians indicated. Each dot represents one mouse. Statistical comparisons were performed using two-tailed Mann–Whitney U tests. Significance: ns (p > 0.05); * (p < 0.05); ** (p < 0.01); *** (p < 0.001); **** (p < 0.0001).

Next, we examined the epitope specificity of the antibody responses induced by immunization. Previous research by our group showed that immunization of rabbits and macaques with ConM Env primarily targets the V1V2 region on the trimer apex as the primary site for neutralizing antibodies *(Sliepen et al. 2019; Reiss et al*., *under revision)*. To determine whether immunization with ConM Env or ConM Env-mC3d, adjuvanted with AddaS03, similarly induces V1V2-directed Ab responses, we tested serum binding to a modified ConM Env trimer in which the V1V2 domain was replaced with that of BG505 SOSIP.664 (ConM Env V1V2 switch). At week 8, the ConM Env-mC3d group exhibited a more V1V2-focused antibody response than the ConM Env group (Fig. 4D; 1.9 vs. 1.0; p < 0.0001). This trend persisted after three immunizations, with the ConM Env-specific response in the ConM Env-mC3d group remaining significantly enriched for V1V2-directed antibodies compared to the ConM Env group (1.5 vs. 1.1; p = 0.007). However, neither group reached the >2.5 ratio threshold (set as the criterion for a robust dominant V1V2-specific response) at either time point (Fig. 4D), suggesting that while V1V2 targeting was enhanced by C3d fusion, the overall magnitude of this response remained modest.

To further characterize the specificity of the antibody response, we assessed recognition of the trimer base using a modified ConM Env construct (ConM Env-PAS50) in which a fifty–amino acid proline–alanine–serine (PAS; *Schlapschy et al. 2013; Kraft et al. 2022*) extension was fused to the C-terminus to sterically mask base epitopes. At week 8, mice immunized with ConM Env-mC3d showed significantly reduced base-specific antibody responses compared to the ConM Env group (Fig. 4E; 1.0 vs. 1.6; p < 0.0001). This pattern was maintained at the final bleed, with lower binding ratios in the ConM Env-mC3d group (0.9 vs. 1.6; p < 0.0001). As with the V1V2 switch, differential binding ratios in both groups remained above the <0.4 threshold, indicating that base reactivity, while reduced in the C3d group, was still detectable. Nonetheless, the combination of increased V1V2-directed responses and reduced base-specific antibodies in the ConM Env-mC3d group suggests that C3d fusion helps focus the early antibody response towards more desirable epitopes.

Consistent with the observed epitope focusing, the ConM Env-mC3d group demonstrated significantly enhanced ConM virus neutralization at both week 8 and the final bleed compared to the unconjugated ConM Env trimer with AddaS03 (p = 0.0003 and p < 0.0001, respectively; Figs. 4F–G), despite similar levels of total ConM Env-specific antibody titers after three immunizations. To further assess the breadth of the neutralizing antibody response, we tested serum samples against the more neutralization-resistant but closely related ConS pseudovirus. Although most mouse sera failed to neutralize ConS effectively, sporadic low-level activity was detected. At the final bleed, mice immunized with ConM Env-mC3d plus AddaS03 showed a trend toward enhanced ConS neutralization compared to the other groups (Fig. 4H–I; p = 0.052). These findings suggest that while the effects of C3d fusion are most pronounced early in the immunization schedule, the resulting antibody response remains more focused on desirable epitopes, potentially contributing to the significantly enhanced neutralization observed after three immunizations.

### C3d Fusion Promotes Differentiation of ConM Env-specific B cells into Classed-Switched Memory B cells

To evaluate the impact of C3d fusion on memory B cell differentiation, we analyzed splenocytes collected four weeks after the final immunization from the same male mice (N=5 per group) immunized with ConM Env or ConM Env-mC3d at weeks 0, 4, and 8 with AddaS03, as described above. We observed no significant differences between groups in the total number of CD19^+^ B cells or total memory B cells, including both unswitched and class-switched memory populations (Fig. S3A, Figs. S3C-D). Notably, we observed a higher frequency of GC B cells in ConM Env-mC3d–immunized mice compared to ConM Env (6.3% vs. 3.2%, p = 0.0317; Fig. S3B), although total ConM Env-specific B cells and ConM Env-specific GC B cells remained similar between groups after three immunizations (Figs. S3E-F).

Phenotypic analysis of ConM Env-specific memory B cells revealed that ConM Env immunization resulted in a higher percentage of unswitched memory B cells compared to ConM Env-mC3d (0.004% vs. 0.001%, p = 0.016; Fig. S3G). In contrast, ConM Env-mC3d immunization led to a higher proportion of class-switched ConM Env-specific memory B cells compared to ConM Env alone (0.6% vs. 0.4%, p = 0.008; Fig. S3H), indicating a more differentiated memory B cell response. Together, these findings suggest that while overall Env-specific B cell numbers are similar after three immunizations, C3d fusion enhances the quality of the memory B cell response by promoting class-switching, supporting the generation of a more functionally mature memory compartment.

### Enhanced Formation of ConM Env-specific B Cells and Germinal Center B Cells in Active Germinal Center Reactions

Since the differences between ConM Env and ConM Env-mC3d diminished after three immunizations, apart from improved neutralization, we sought to understand the underlying mechanisms by examining early GC dynamics during ongoing GC reactions. To explore this, we immunized female and male mice with fluorescently labeled ConM Env or ConM Env-mC3d, using AddaS03 as an adjuvant and tracked GC B cell formation over time (Fig. 5A; Fig. S4A). As expected, initial phenotyping at 24 hours post-immunization, serving as a baseline, revealed no differences in GC B cell formation (Fig. S4A; identified as CD38^-^, CD138^-^, GL7^+^, and CD95^+^ B cells) between ConM Env and ConM Env-mC3d (Fig. 5B). By day 7, however, GC B cell formation was significantly enhanced in mice immunized with ConM-Env-mC3d compared to those receiving ConM Env alone (Fig. 5B; p = 0.002), indicating that C3d fusion promotes early GC dynamics.

**Figure 5:**
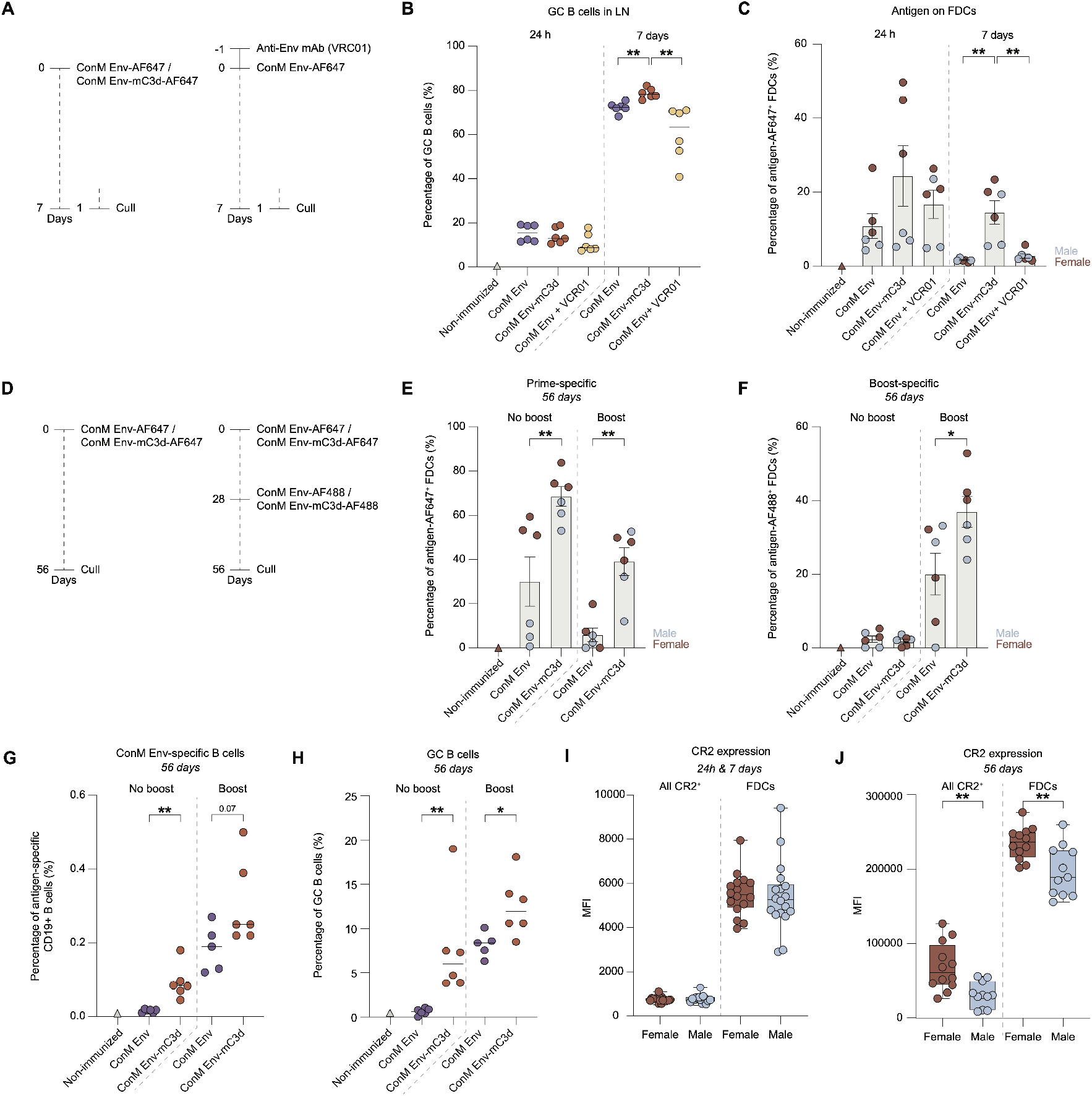
Antigen capture, retention, and GC B cell responses in draining lymph nodes following ConM Env and ConM Env-mC3d immunization. **.(A)** Immunization schedule of mice for analyzing antigen localization in draining lymph nodes (LNs) after immunization with ConM Env-AF647 and ConM Env-mC3d-AF647, each supplemented with AddaS03 adjuvant. Six mice per group were used, split evenly between males and females (3 each). Left side depicts the timeline for evaluating antigen presence at 24 hours and 7 days post-injection. On the right, a similar setup is used, with the addition of VRC01, an anti-HIV-1 Env-specific monoclonal antibody (mAb), injected intraperitoneally (i.p.) one day prior to subcutaneous administration of AF647-labeled antigen to assess formation and localization of antigen-immune complexes. **(B)** Representative percentage of live CD19^+^ CD95^+^ GL7^+^ GC B cells in lymph nodes from mice treated as described in Figure 5A, assessed at 24 h and 7 days post-immunization with ConM Env-AF647, ConM Env-mC3d-AF647, or ConM Env-AF647 with prior i.p. administration of VRC01 (ConM Env + VRC01). **(C)** Percentage of antigen-AF647^+^ FDCs (CD45^-^,B220^-^,CD31^-^,PDPN^+^,CD21/35^hi^,Madcam1^+^) at 24 hours and 7 days post-immunization in mice treated with ConM Env-AF647, ConM Env-mC3d-AF647, and ConM Env-AF647 + VRC01. Sex differentiation is shown with females in maroon and males in light blue. **(D)** Immunization schedule of mice for assessing long-term antigen retention and effect of GCs in draining lymph nodes. Left: mice were immunized with ConM Env-AF647 or ConM Env-mC3d-AF647. Right: mice were immunized with ConM Env-AF647 or ConM Env-mC3d-AF647 and boosted on day 28 with their AF488-labeled counterparts. All immunizations included AddaS03 adjuvant. Mice were culled at day 56 (n=6 per group, 3 male and 3 female). **(E)** Percentage of total antigen-AF647^+^ FDCs (prime-specific) for mice treated as described in Figure 5D, assessed at 56 days post-immunization. Sex differentiation is shown with females in maroon and males in light blue. **(F)** Percentage of total antigen-AF488^+^ FDCs (boost-specific) for mice treated as described in Figure 5D, assessed at 56 days post-immunization. Sex differentiation is shown with females in maroon and males in light blue. **(G)** Percentage of ConM Env-specific CD19^+^ B cells assessed at 56 days post-immunization, for mice treated as described in Figure 5D. **(H)** Percentage of CD19^+^ CD95^+^ GL7^+^ GC B cells for mice treated as described in Figure 5D. **(I)** Median fluorescence intensity (MFI) of CR2/CD21 expression in female and male mice immunized as in Figure 5A, measured on all CR2^+^ cells and on FDCs (CD21^hi^ Madcam1^+^) from lymph nodes at 24 h and 7 days post-immunization (pooled). **(J)** Median fluorescence intensity (MFI) of CR2/CD21 expression in female and male mice immunized as in Figure 5D, measured on all CR2^+^ cells and on FDCs (CD21^hi^ Madcam1^+^) from lymph nodes at 56 days post-immunization. Data are shown as mean ± SEM, median with scatter, or box-and-whisker plots (min to max), as indicated. Each dot represents one mouse. Statistical comparisons were performed using two-tailed Mann–Whitney U tests, except in panels I and J (unpaired t-tests). Significance: ns (p > 0.05); * (p < 0.05); ** (p < 0.01); *** (p < 0.001).

To further examine how C3d fusion influences early GC dynamics, we compared it to immune complex (IC) formation—a process known to impact GC responses. To generate ICs *in vivo*, mice received an intravenous injection of the anti-Env monoclonal antibody VRC01 one day prior to immunization with ConM Env (Fig. 5A). At 24 hours post-immunization, GC B cell frequencies were again similar between the ConM Env-mC3d and ConM Env-IC groups (13 vs 8.4%; Fig. 5B). By day 7, however, mice immunized with ConM Env-mC3d exhibited significantly higher levels of GC B cells than those in the ConM Env-IC group (p = 0.002), suggesting that C3d fusion effectively promotes early GC expansion, even in the presence of antibody–antigen immune complexes.

We next examined the effect of C3d on the longevity of the GC reaction (Fig. 5D). At 56 days post-immunization, without additional boosting, the lymph nodes of the ConM-Env-mC3d group showed significantly increased GC B cell (6.0 vs 0.6%, p = 0.002; Fig. 5E) and ConM Env-specific B cell formation (0.084 vs 0.0175%, p = 0.002; Fig. 5G) compared to ConM Env, underscoring the potential of C3d fusion to enhance antigen-specific germinal center formation and boost immunogenicity. Similar results were observed after receiving a boost, although the effects became less pronounced (Figs. 5G-H). These findings demonstrate the potential of C3d fusion to Env immunogens to improve vaccine efficacy by strengthening early GC responses and promoting antigen-specific B cell development.

### Enhanced Antigen Retention and Longevity of ConM Env-mC3d in Germinal Centers

Given the enhanced GC responses observed with ConM Env-mC3d, we next examined whether C3d fusion influences antigen retention and persistence on FDCs within GCs. To do so, we immunized mice with fluorescently labeled ConM Env or ConM Env-mC3d, using AddaS03 as an adjuvant (Fig. 5A), and analyzed antigen deposition on FDCs in draining lymph nodes using flow cytometry at 24 hours, followed by assessments of antigen retention at 7 and 56 days post-immunization (Fig. S4B). We observed substantial heterogeneity in antigen deposition between female and male mice, with females showing higher antigen deposition/presentation on FDCs despite similar overall FDC frequencies across groups and all timepoints (Figs. 5C, 5E, and Fig. S4C). At 24 hours post-immunization, FDC-associated antigen levels were not statistically different across groups when sexes were pooled, though ConM Env-mC3d showed a 2.25-fold increase over ConM Env (Fig. 5C). However, ConM Env-mC3d induced significantly higher FDC loading in female mice compared to ConM Env (p = 0.031) and ConM-IC (p = 0.029), indicating enhanced antigen capture in females (Fig. S4C). By seven days post-immunization, antigen retention was consistently higher for ConM Env-mC3d compared to ConM Env or ConM-IC for both sexes (p = 0.0038 and p = 0.0043, respectively; Fig. 5C).

Analysis at day 56 revealed that the enhancement in antigen retention with C3d fusion was durable and persisted over time (Fig. 5D-E). By day 56 post-immunization, ConM Env-mC3d continued to exhibit significantly higher levels of FDC-bound antigen compared to ConM Env (66% vs. 30%, p = 0.009; Fig. 5E), indicating that C3d fusion promotes prolonged antigen persistence on FDCs. Analysis at day 56 also showed that additional boosting at day 28 (Fig. 5A) reduced the retention of the primary antigen on FDCs (Fig. 5E, right), likely due to replacement or competition from the newly introduced boost antigen. In addition, 39% of the FDCs from ConM Env-mC3d-immunized mice retained the priming antigen (AF647) and 37% the boost antigen (AF488), whereas only 5.9% and 20% of FDCs, respectively, from ConM Env-immunized mice retained the priming and boost antigens (Fig. 5E-F). These differences were statistically significant for both prime (p = 0.004) and boost (p = 0.04) antigen retention at day 56, indicating that C3d fusion enhances both the durability and cumulative presentation of antigen on FDCs following booster immunization.

Interestingly, this prolonged antigen presentation in females coincided with higher CR2 expression at day 56, both in total CR2^+^ cells and specifically on FDCs (Fig. 5J). No sex-specific differences in CR2 expression were observed at earlier time points (Fig. 5I), suggesting that enhanced long-term antigen retention in females may be partly explained by increased CR2 availability during later stages of the response. Consistent with this, CR2 expression levels moderately correlated with antigen levels retained on FDCs 56 days post-immunization (Pearson’s R^2^ = 0.49 and p = 0.017; Fig. S4E).

These findings highlight the potential of C3d fusion to ConM Env in enhancing initial immune responses and sustaining antigen retention in GCs over time and through boosting, which could improve vaccine efficacy through prolonged antigen availability.

## Discussion

Developing HIV-1 vaccines capable of eliciting durable and bNAb responses remains a major challenge, owing to the poor immunogenicity of Env glycoproteins and the difficulty in sustaining GC activity. In this study, we demonstrate that fusing the complement fragment C3d to the ConM SOSIP trimer robustly enhances multiple facets of the early GC response. Specifically, the C3d fusion improved B cell activation, promoted more efficient antigen display by FDCs, and enhanced both the formation and persistence of GC B cells *in vivo*. These immunological enhancements were accompanied by increased antibody titers, refined epitope targeting, and improved autologous virus neutralization, suggesting that C3d engages complementary mechanisms—B cell co-stimulation and prolonged antigen retention—to shape early GC dynamics. Collectively, these findings establish C3d fusion as a promising strategy for improving Env-based vaccines and sustaining humoral immunity.

Efforts to elicit protective antibodies against HIV-1 have been hindered by the dense glycan shield of the Env trimer *(Berndsen et al. 2020; Stewart-Jones et al. 2016)*, which obscures conserved neutralizing sites and redirects the antibody response toward more accessible, immunodominant but non-neutralizing regions such as the trimer base — a neo-epitope inaccessible on membrane-bound Env but exposed on soluble trimers *(Antanasijevic et al. 2021; Cottrell et al. 2020; Kwong et al. 2002)*. In this study, we found that fusing C3d to the ConM Env trimer promoted more focused antibody targeting. Notably, immunization with ConM Env-mC3d reduced antibody reactivity to base epitopes, while increasing recognition of the V1V2 apex — a key neutralization determinant *(Sliepen et al. 2019; Reiss et al*., *under revision)*. This shift in epitope targeting was supported by ELISA data showing that RM19R, a base-directed antibody, failed to bind the C3d-fused trimer, suggesting that C3d may sterically hinder access to the base. This redirection of the antibody response was accompanied by improved autologous virus neutralization, indicating that occlusion of non-neutralizing epitopes may help steer the immune response toward more protective antibody specificities.

C3d fusion significantly enhanced early B cell activation and GC responses, likely contributing to the improved magnitude and quality of the antibody response. Consistent with previous work demonstrating the ability of C3d to lower the activation threshold of B cells through CR2 signaling *(Dempsey et al. 1996; Bale et al. 2023)*, we observed stronger B cell activation even at lower antigen doses. This early enhancement was accompanied by more robust GC responses, which likely supported the observed improvements in epitope targeting and neutralization, as GC dynamics play a central role in shaping antibody specificity and affinity. Furthermore, we detected a higher frequency of class-switched, ConM-specific memory B cells in the ConM Env-C3d group, whereas immunization with ConM Env alone yielded more non-switched memory B cells, suggesting that C3d also promotes more functionally mature memory B cell differentiation. By promoting early B cell activation and facilitating efficient GC initiation, C3d fusion establishes an environment that favors the maturation of high-affinity, protective antibodies.

Prolonged antigen availability within lymphoid follicles is increasingly recognized as a critical factor for sustaining GC activity and guiding antibody maturation *(Victora and Nussenzweig 2022; Cirelli et al. 2019; Cirelli and Crotty 2017; Tam et al. 2016)*. In our study, C3d fusion significantly improved antigen retention on FDCs, with intact ConM Env-mC3d detectable on FDCs as late as 56 days post-immunization. This enhanced retention was particularly pronounced in female mice at early time points and correlated with a higher frequency of antigen-loaded FDCs compared to animals immunized with ConM Env. Notably, we also demonstrated improved display of C3d-fused Env to HIV-1-specific B cells using human FDCs, underscoring the translational relevance of this mechanism. These findings suggest that C3d acts not only to enhance B cell activation, but also to promote the long-term deposition of antigen in B cell follicles, thereby supporting more sustained GC responses. This strategy mirrors the benefits observed with slow-delivery and dose-escalation regimens *(Aung et al. 2023; Cirelli et al. 2019; Lee et al. 2022; Meany et al. 2025)*, which rely on extended antigen availability to optimize affinity maturation. However, C3d fusion achieves this effect through molecular design rather than complex delivery systems, offering a practical and scalable approach to improving Env immunogenicity.

While C3d fusion significantly enhanced early immune responses, including GC dynamics and antibody titers, these benefits diminished after multiple immunizations, likely due to antibody feedback that suppresses the activation of new B cells and limits the breadth of the response. This has been observed in other studies, where high antibody levels can impair GC activation and affinity maturation *(Cirelli and Crotty 2017; Coelho et al. 2023; Cirelli et al. 2019)*. The reduced differences in B cell phenotype and antibody titers between ConM Env-mC3d and unconjugated ConM Env over time likely reflect the increase in antibody affinity with repeated exposure, activating sufficient GC activity even without further C3d fusion enhancement. Additionally, epitope masking by pre-existing antibodies may have limited the ability of subsequent boosts to further enhance antibody titers to these epitopes *(Zarnitsyna et al. 2016; Schaefer-Babajew et al. 2023; Brown et al. 2025)*. Despite these challenges, the early advantages of C3d fusion highlight its potential as an effective priming strategy, especially for germline-targeting HIV-1 antigens, and support its use in optimized prime-boost strategies to shape long-term immune responses.

An intriguing observation in this study was the sex-based differences in immune responses, with female mice exhibiting stronger antibody responses and higher antigen retention in draining lymph nodes compared to males. This was accompanied by higher CR2 expression in female mice at later time points, which may partly explain the enhanced retention and immunogenicity observed in females. These findings align with previous studies showing that females generally exhibit stronger humoral immune responses, possibly influenced by hormonal regulation *(Potluri et al. 2019; Anticoli et al. 2023; Fischinger et al. 2019)*. Estradiol, the primary estrogen in mice and humans, has been shown to enhance GC B cell responses, increase Tfh cell differentiation, and promote higher antibody production *(Klein and Flanagan 2016; Dhakal et al. 2024)*. Interestingly, a human clinical trial conducted by our group using ConM Env also reported sex-based differences in immune responses *(Reiss et al*., *under revision)*. In that trial, TLR4-driven adjuvant activity was thought to contribute to these differences, but our data suggest that immunization-induced elevation of CR2 expression and adjuvant effects may also play a significant role in enhancing immune responses in females. These findings underscore the importance of considering sex as a biological variable in vaccine studies. Further research into its role in antigen retention, GC dynamics, and immune activation could help refine vaccine strategies for optimal efficacy across sexes.

The combination of AddaS03 and C3d played a crucial role in enhancing the immunogenicity of the immunogens in mice in this study. While C3d alone showed limited efficacy as an adjuvant, its combination with AddaS03 proved synergistic. AddaS03 activates Th1, Th2, and Th17 pathways, providing cytokines and co-stimulatory signals crucial for GC B cell maturation and affinity maturation *(Yam et al. 2015; Neeli et al. 2023)*. It also enhances antigen uptake and dendritic cell maturation, further strengthening adaptive immune responses *(Nakkala et al. 2024)*. Meanwhile, C3d amplifies B cell activation through CR2 engagement and enhances antigen retention on FDCs. Together, these complementary mechanisms drive the robust GC activity and improved neutralization responses observed at early time points. While this study primarily focused on B cell dynamics, the broader impact of AddaS03 and C3d on T cell responses remains under-explored. Investigating how this combination influences long-term immunity could provide valuable insights for optimizing future adjuvant strategies.

Overall, this study demonstrates the potential of C3d fusion as a promising strategy to enhance the germinal center response of HIV-1 Env immunogens. While improved neutralization is a key outcome, our findings reveal that C3d achieves this by fundamentally shaping GC dynamics and extending antigen retention on FDCs—mechanisms that are often overlooked in vaccine studies. By sustaining antigen availability and promoting early B cell activation, C3d fusion provides a valuable framework for optimizing vaccine design, particularly for challenging immunogens such as HIV-1 Env, where robust B cell engagement and prolonged GC activity are critical for vaccine efficacy.

## Materials and Methods

### Design of ConM Env constructs

The ConM SOSIP version 7 gp140 (ConM Env) plasmid was generated as previously described *(Sliepen et al. 2019)*. For the ConM Env-C3d constructs, a sequence was designed with a 12-amino acid GSG linker, followed by either human or murine C3d (Integrated DNA Technologies). The ConM Env v7 plasmid was digested using *BamHI* and *NotI* Fast-digest enzymes and assembled with the C3d sequence by Gibson Assembly, yielding ConM Env-hC3d and ConM Env-mC3d constructs.

To generate the ConM Env-PAS50 construct, we designed a codon-optimized DNA fragment encoding a short GSGG linker followed by a 50–amino acid pro-line–alanine–serine (“PAS50; *Schlapschy et al. 2013)* The PAS50 amino acid sequence is: GSGGSASPAAPAPASPA APAPSAPAASPAAPAPASPAAPAPSAPAASPAAPAPA SPAAPAPSAPAASPAAPAPASPAAPAPSAPAASPAAP APASPAAPAPSAPAASPAAPAPASPAAPAPSAPAASP AAPAPASPAAPAPSAPAASPAAPAPASPAAPAPSAPA ASPAAPAPASPAAPAPSAPA.This fragment was synthesized (Integrated DNA Technologies) and inserted into the *BamHI/NotI*-digested ConM Env v7 plasmid using Gibson Assembly. All constructs were verified by DNA sequencing prior to protein expression.

### Prediction of ConM Env-C3d structures

The amino acid sequence of ConM Env-C3d fusion constructs was used to generate a prediction using an open-access AlphaFold3 server (alphafoldserver.com). The resulting predictions were analyzed and visualized using ChimeraX-1.9 *(Pettersen et al. 2021; Meng et al. 2023)*.

### Protein expression and purification

All protein antigens—ConM Env, ConM Env-mC3d, ConM Env-hC3d, ConM-BG505 SOSIP V1V2 variant, and ConM Env PAS50 proteins—were expressed and purified as previously described *(Sliepen et al. 2019)*. Briefly, HEK293F cells (Invitrogen, cat# R79009) were transiently transfected with a 4:1 ratio of Env plasmid to furin expression plasmid to promote optimal furin-cleavage of SOSIP constructs. Cultures were maintained for seven days, after which the supernatants were harvested and vacuum-filtered using 0.22 µm Steritops (Merck Millipore). Env proteins were purified via affinity chromatography using CNBr-activated Sepharose 4B resin (GE Healthcare) conjugated to the PGT145 broadly neutralizing antibody. Bound proteins were eluted with 3 M Mg_2_Cl_2_ pH 7.8, directly into a neutralization buffer (20 mM TrisHCl pH 8.0, 75 mM NaCl). To purify only trimeric forms, eluted proteins were subjected to size exclusion chromatography on a Superdex 200 10/330 G/L column (GE Healthcare) only. The trimeric fractions were then concentrated and buffer-exchanged into PBS using 100 kDa VivaSpin20 centrifugal filters. Final protein concentrations were determined using a NanoDrop 2000 spectrophotometer (Thermo Fisher Scientific).

### ELISA

Galanthus nivalis lectin (Vector Laboratories) was coated onto half-area 96-well plates at a concentration of 20 µg/mL in 0.1 M NaHCO_3_ (pH 8.6) and incubated overnight at room temperature (RT). Plates were washed with TBS and blocked for 30 min using Casein blocking buffer (Thermo Scientific). Purified Env proteins (2 µg/mL) were added and incubated for 2 hours at RT. After three washes with TBS, monoclonal antibodies (PGT145, PGT121, PGT151, PGDM1400, VRC01, 2G12, RM19R, and COVA1-16; 1 µg/mL) or mouse CR2 protein (2 µg/mL; Enquire BioReagents) were added and serially diluted (1:3). Plates were incubated for an additional 2 hours at RT. Following three washes, HRP-conjugated donkey anti-human IgG (Jackson ImmunoResearch) was added at a 1:3000 dilution in Casein buffer and incubated for 1 hour at RT. Plates were washed five times with 1× TBS containing 0.05% Tween-20. Color development was initiated by adding freshly prepared substrate solution (1% 3,3’,5,5’-tetramethylbenzidine, 0.01% hydrogen peroxide, 0.1 M sodium acetate, 0.1 M citric acid) and allowed to proceed for 2 min. The reaction was stopped with 0.8 M H_2_SO_4_, and absorbance was measured at 450 nm (OD_450_).

### Generation of HIV-specific B cells

HIV-specific Ramos B cells were produced as previously described *(Brouwer et al. 2021)*. In short, PGDM1400 and PGT121 HIV-specific Ramos B cells were generated by replacing the immunoglobulin variable regions of the pRRL EuB29 2-1261gl IgG TM.BCR.GFP.WPRE plasmid with those from the PGDM1400 or PGT121 antibodies using Gibson assembly (Integrated DNA Technologies). Lentivirus was produced by co-transfecting HEK293T cells with the newly generated expression plasmid together with the pVSV-g, pMDL, and pRSV-Rev plasmids using lipofectamine 2000 (Invitrogen). Two days after transfection, IgM-negative Ramos B cells were transduced using the harvested HEK293T supernatant. Seven days following infection, B cells double-positive for IgG and GFP were sorted using a FACS Aria-II SORP (BD Biosciences) and were subsequently expanded and maintained in long-term culture.

### Calcium flux assay

B cell activation experiments of the generated HIV-specific B cells were performed as previously described *(Brouwer et al. 2021)*. PGDM1400 or PGT121 Ramos B cells (4 million/ml) were pre-stained with 1.5 µM Indo-1 (Invitrogen), washed and re-suspended in HBSS supplemented with calcium and magnesium (Gibco) and 0.5% FCS at 1 million cells/ml. PGT121 and PGDM1400 B cell Ca^2+^ influx upon antigen stimulation was assessed by measuring Indo-1 fluorescence (379/450 nm emission ratio) upon UV excitation on a LSR Fortessa flow cytometer (BD Biosciences). Following 30s of baseline measurement, aliquots of 1 million cells/mL were then stimulated for 500 s at RT with either ConM, ConM Env-hC3d, or ConM with human complement C3d (hC3d; Prospec). For the stimulation, ConM and hC3d were added in an equimolar ratio with ConM Env-hC3d. Maximum Indo-1 fluorescence was established using 1 mg/mL ionomycin (Invitrogen). Kinetics analyses were performed using FlowJo v10.10.

### Human FDC isolation

Human follicular dendritic cells (FDCs) were isolated from anonymously donated palatine tonsils collected during routine tonsillectomies at the Diakonessenhuis Utrecht, following a modified version of the protocol by *Breeuwsma and Heesters 2023*. Tonsils were rinsed with ice-cold PBS supplemented with penicillin/streptomycin (P/S), mechanically dissociated using sterile scissors and a metal sieve, and collected in ice-cold PBS. Tissue fragments were briefly centrifuged and enzymatically digested in HBSS containing 20 mM HEPES, 0.5% FCS, 200 U/mL collagenase IV, 6 U/mL Dispase II, and 50 µg/mL DNase I for 1 hour at 37°C with gentle rocking. Following digestion, cells were washed in sorting buffer (HBSS without calcium/magnesium, 20 mM HEPES, 5% FCS, 2 mM EDTA, and gentamicin) and filtered through a fine metal sieve. Mononuclear cells were enriched by discontinuous Percoll™ gradient centrifugation (21% and 40%) at 300 x g for 25 min at room temperature. The FDC-containing interphase was collected, washed, and incubated with CD35-biotin, CD11a, and CD49d antibodies (BioLegend), followed by magnetic separation using streptavidin-conjugated beads and the MojoSort™ system (BioLegend). Isolated FDCs were cultured in IMDM supplemented with HEPES, non-essential amino acids, sodium pyruvate, 5% FCS, and 100 I/mL p/s in plates pre-coated with poly-D-lysine (100 µg/mL), laminin (40 µg/mL), and type I collagen (100 µg/mL).

### Antigen presentation by FDCs to human HIV-specific B cells

#### (i) Antigen Labeling and Immune Complex Formation

ConM Env, ConM Env-C3d, and SARS-CoV-2 Wuhan spike (control) proteins were fluorescently labeled using an AF647 microscale protein labeling kit, following the manufacturer’s instructions (Fisher Scientific). Epitope accessibility was confirmed after labeling. Immune complexes (ICs) were formed by incubating a 1:2.5 ratio of protein to anti-HIV-1 Env 2G12 antibody with or without 10% human serum (Sigma) in GVB++ buffer (in-house Gelatin Veronal Buffer with Ca^2+^ and Mg^2+^) for 30 min at 37 °C. The protocol IC formation and FDC loading was adapted from Heesters et al. (2021) *(Heesters et al. 2021)*.

#### (ii) Antigen Uptake and Surface Retention by FDCs

Labeled ICs or recombinant proteins were added to cultured human FDCs for 1 hour at 37 °C. Cells (20,000 per well) were washed three times with pre-warmed media to remove unbound antigen, stained in the dark for 30 min at 37 °C in FACS buffer (PBS + 1 mM EDTA + 2% FCS) containing a viability dye (Invitrogen), PE-Cy7 anti-human PDPN (Clone: NC-08; Biolegend), PE anti-human CD35 (Clone: E11, Biolegend), BUV395 anti-human CD21 (Clone: B-ly4; BD Optibuild) then washed and analyzed on a BD LSR Fortessa. Data was processed using FlowJo v10.8.

#### (iii) Antigen Presentation to HIV-Specific B Cells

Following antigen uptake (as described above), FDCs were washed three times with pre-warmed FDC medium and incubated with 20,000 PGT121 Ramos B cells per well for 3 hours at 37 °C. After co-culture, B cells were harvested and stained in FACS buffer (PBS + 1 mM EDTA + 2% FCS) with viability dye-eF780 (Invitrogen) and CD19-BUV496 (Biolegend). Stained cells were washed and then analyzed by flow cytometry using a BD LSR Fortessa, and data were processed with FlowJo v10.8. Antigen uptake by PGT121 B cells was quantified by gating on live CD19^+^ GFP^+^ antigen-AF647^+^ lymphocytes.

### Antigen retention in GCs by FDCs

#### (i) Immunization

Isoflurane-anesthetized C57BL6 mice were immunized subcutaneously (s.c.) with equimolar concentrations of ConM Env-AF647 (10 µg/injection) or ConM Env-C3d-AF647 (15.83 µg/injection) and 50 µl of AddaS03 in a total volume of 100 µl per injection. Each animal received four injections, one in each arm and leg pit. At endpoint (24 h, 7 days, or 56 days post-immunization), the axillary, brachial, and inguinal lymph nodes were collected and analyzed. When specified, mice underwent a booster immunization on day 28 (ConM Env-AF488 or ConM Env-C3d-AF488 in AddaS03) and tissues were collected on day 56. In the case of VRC01 treatment groups, mice received an intraperitoneal (i.p.) injection of 100 µg VRC01 antibody. After 24 hours, mice were s.c. injected with 10 µg of ConM-AF647 as described above, and tissues were collected either 24 hours or 7 days post-immunization.

#### (iii) FDC isolation and immunophenotyping

Draining LNs were mechanically disaggregated into small pieces. Dissociated tissue was then incubated in digestion buffer (RPMI-1640 medium containing 2% FCS, 20 mM HEPES pH7.2, 0.1 mg/ml collagenase P and 25 µg/ml DNase I) at 37°C for 60 min (15 min x 4 times). After digestion, cell suspensions were filtered (100 µm strainer) and washed with FACS buffer (0.5% FCS and 10 mM EDTA in PBS). Single-cell suspensions were incubated with: Fc-block (anti-CD16/32), Fixable Viability Dye-e506 (Invitrogen), B220 (clone RA3-6B2, Biolegend), CD45.2 (104, Biolegend), CD21/35 (7E9, Biolegend), PDPN (8.1.1, Biolegend), Madcam1 (MECA-367, BD Biosciences), CD31 (390, Biolegend) in FACS buffer for 20 min on ice. Cells were analyzed using an LSRFortessa or Aurora flow cytometer and analyzed using FlowJo. FDCs were gated as shown in *Martinez-Riano et al*., *2023*.

### Mouse immunizations

All animal procedures were carried out in accordance with protocols approved by the Emory University School of Medicine Institutional Animal Care and Use Committee. C57BL/6 mice (8 to 12 weeks old; Charles River Laboratories) received subcutaneous immunizations at the base of the tail with either 10 µg of ConM or 15.83 µg of ConM Env-C3d (equivalent to 10 µg of ConM), with or without 50 µl of AddaS03™ (InvivoGen). The total injection volume was adjusted to 100 µl using sterile 1× PBS. Immunizations were given at weeks 0, 4, and 8. Blood samples were collected via cheek bleed at defined intervals after each immunization to collect serum. At week 12, blood and spleens were collected. Spleens were mechanically processed using a 3 ml syringe plunger (BD Biosciences) and passed through 100 µm strainers (Corning) to obtain single-cell suspensions. Splenocytes were treated with ACK lysis buffer (Gibco) to remove red blood cells, washed with 1× PBS, and stored in the vapor phase of liquid nitrogen (−196 °C). Serum samples were used to measure antibody responses using luminex immonoassays and pseudo virus neutralization assays. Freshly isolated lymph node cells were used to examine germinal center T follicular helper (GC-TFH) and B cell (GC-BC) responses by flow cytometry. Splenocytes were used to assess antigen-specific B cell responses using flow cytometry.

### Flow cytometry of (antigen-specific) mouse splenocytes

Fluorescent probes were prepared as previously described *(Brouwer et al. 2020)*. Briefly, biotinylated ConM Env proteins were multimerized with streptavidin conjugated to Alexa Fluor 647 (Biolegend) or Brilliant™ UltraViolet 737 (BUV737; BD biosciences) by mixing at a 2:1 molar ratio (protein:fluorochrome) and incubated for 1 hour at 4 °C. To remove any unbound streptavidin, 10 µM biotin (Genecopoiea) was added and incubated for at least 10 min. The labeled probes were then combined in equal molar amounts to a final concentration of 45.5 pM. Mouse splenocytes were washed twice with FACS buffer (0.5% FCS, 10 mM EDTA in PBS) and pre-incubated on ice for 30 min with fluorescently labeled probes and Fc-block (anti-CD16/32; Biolegend). Following this, an antibody mixture containing GL7-PerCP-Cy5.5 (GL.7; Biolegend), a dump channel (eF780-APC) with CD11c (N418; Invitrogen), CD11b (M1/70; Invitrogen), CD3ε (145-2C11; Invitrogen), and CD49b (DX5; Invitrogen), CD19-R718 (1D3; BD Biosciences), IgD-BV785 (11-26c.2a; Biolegend), CD95-BV605 (SA367H8; Biolegend), CD38-BV421 (90; Biolegend), and CD138-PE-Cy7 (281-2; Biolegend) in FACS buffer was added to the cells with the Fc-block and probe mixture, and cells were incubated for an additional 30 min on ice. Cells were than washed twice in FACS buffer and analyzed on a BD LSR Fortessa. Data was processed using FlowJo v10.8.

### Multiplex assay of mouse serum

The humoral response in mouse serum was measured using a previously described in-house Luminex immunoassay *(Brouwer et al. 2021)*. Briefly, ConM Env v7, ConM Env-mC3d, ConM-BG505 SOSIP V1V2 variant, ConM Env PAS50, recombinant murine C3d, and SARS-CoV-2 Wuhan spike protein (75 µg each, equimolar corrected) were covalently coupled to MagPlex beads (12.5 million beads per protein) using a two-step carbodiimide reaction. Optimized dilutions (1:50,000 and 1:1,500,000) of serum samples were incubated with the beads overnight, followed by detection with goat anti-mouse IgG-PE (Southern Biotech). The Magpix platform (Luminex) was used to measure mean fluorescence intensity (MFI), representing the median signal from ~50 beads per well. Background fluorescence was subtracted using MFI values from buffer and beads-only controls.

### IgG Isolation from mouse serum

Mouse IgG was purified from serum using a Protein G Sepharose Fast Flow column (300 µL, Sigma-Aldrich) for subsequent neutralization assays. Briefly, the 100-fold diluted serum diluted in PBS was loaded onto the equilibrated column. After washing with PBS, mouse IgG was eluted with 0.1 M glycine, pH 2.5 (Thermo Fisher Scientific) directly into a neutralization buffer (2 M Tris, pH 8) to prevent protein denaturation. The neutralized eluate was then concentrated and washed using a Vivaspin 6 (100 kDa MWCO, Sartorius). The final purified IgG was filter-sterilized with a 0.22 µm membrane (Corning) and its presence confirmed by Nanodrop (Thermo Scientific) at 280 nm.

### ConM and ConS virus neutralization assay

The serum neutralization capacity was assessed using the TZM-bl neutralization assay, as previously described (Montefiori 2009). TZM-bl cells were seeded one day prior in DMEM supplemented with 10% FCS and 100 U/mL P/S, aiming for 70–80% confluency at assay time. IgG purified from mouse serum was heat-inactivated (56 °C, 60 min, 1:2 in PBS), diluted 1:50 in culture medium, and serially diluted in triplicate. Dilutions were mixed 1:1 with either ConM virus or ConS pseudovirus and incubated at room temperature (60 min for ConM, 30 min for ConS). Mixtures were added 1:1 to TZM-bl cells preincubated with medium containing 40 µg/mL DEAE-dextran and 400 nM saquinavir. After 72 h at 37 °C, cells were lysed, and luciferase activity was measured using Bright-Glo reagent (Promega) on a Glomax plate reader. RLUs were normalized to virus-only controls; media-only wells served as background. VRC01 was included as a positive control to validate assay performance. Neutralization titers (ID50) were defined as the IgG dilution at which infectivity was reduced by 50%, based on a non-linear regression fit in GraphPad Prism (version 10.5.0).

### Statistical analysis

All statistical analyses were performed using GraphPad Prism (version 10.5.0). Statistical tests were chosen based on data distribution, sample size, and experimental design. Two-tailed Mann–Whitney U tests were used for unpaired comparisons when normality could not be assumed. Unpaired t-tests were applied to normally distributed unpaired data, and paired t-tests were used for matched comparisons (e.g., human donor assays). Normality of data distribution was assessed using the Shapiro–Wilk test. Correlations were assessed using Pearson correlation coefficients, applied under the assumption of normally distributed variables. Significance was defined as p < 0.05 and is indicated as follows: ns (p > 0.05); * (p < 0.05); ** (p < 0.01); *** (p < 0.001); **** (p < 0.0001).

## Acknowledgments

We thank Drs. Li Wu and Vineet N. Kewal Ramani from the NIH AIDS Reagent Program for kindly providing Ramos B cells, and Dr. Andrew McGuire for the pRRL.EuB29 lentiviral vector used to transduce these cells. We are also grateful to Ilja Bontjer and Kwinten Sliepen for their development of the ConM Env immunogen. Some figures were created with BioRender.com. We would like to thank the animal staff in the rodent vivarium at the Emory National Primate Research Center (ENPRC) for assistance with animal care in this study.

## Supplementary Information

**Figure S1:**
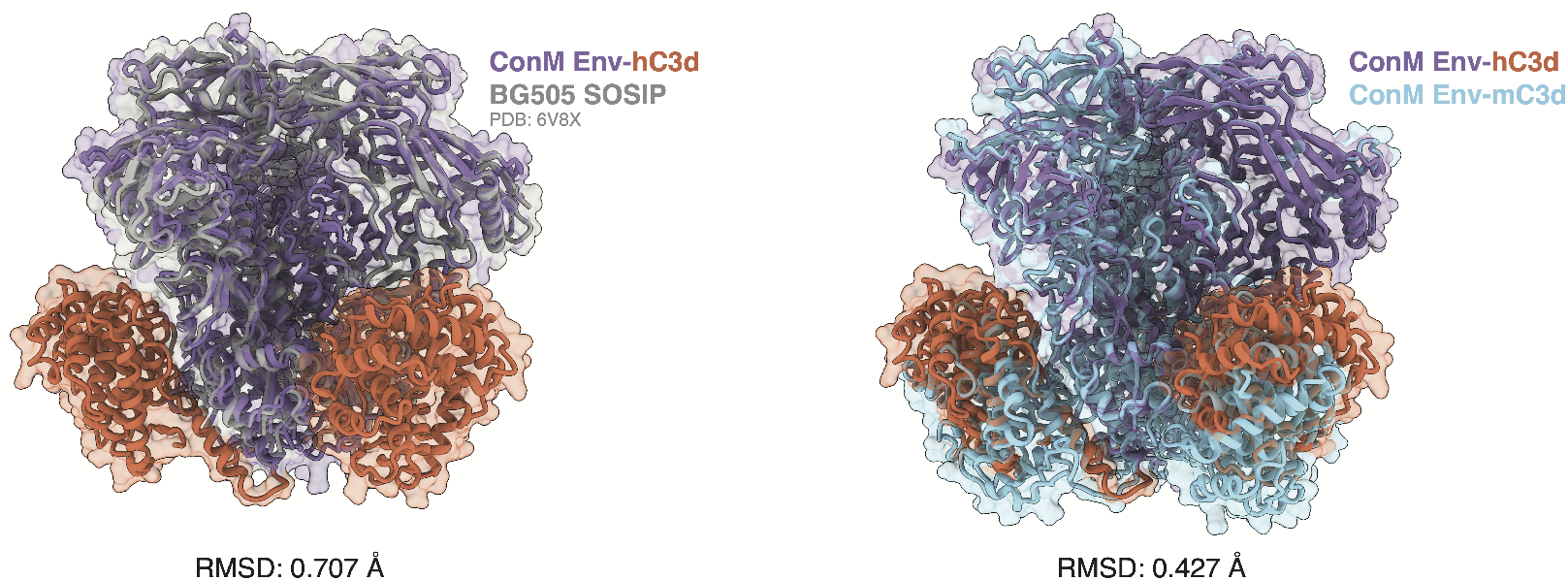
Structural comparison of ConM Env-hC3d with BG505 SOSIP and ConM Env-mC3d. Comparison of predicted ConM Env-hC3d with BG505 SOSIP (left) and ConM Env-murine C3d (right). BG505 SOSIP (PDB: 6V8X, left panel) and AlphaFold3-predicted ConM Env-mC3d show a high degree of similarity with the predicted ConM Env-hC3d structure. Root mean square deviation (RMSD) values are a measure of the average distance between atoms of superimposed structures, with lower values indicating a better fit.

**Figure S2:**
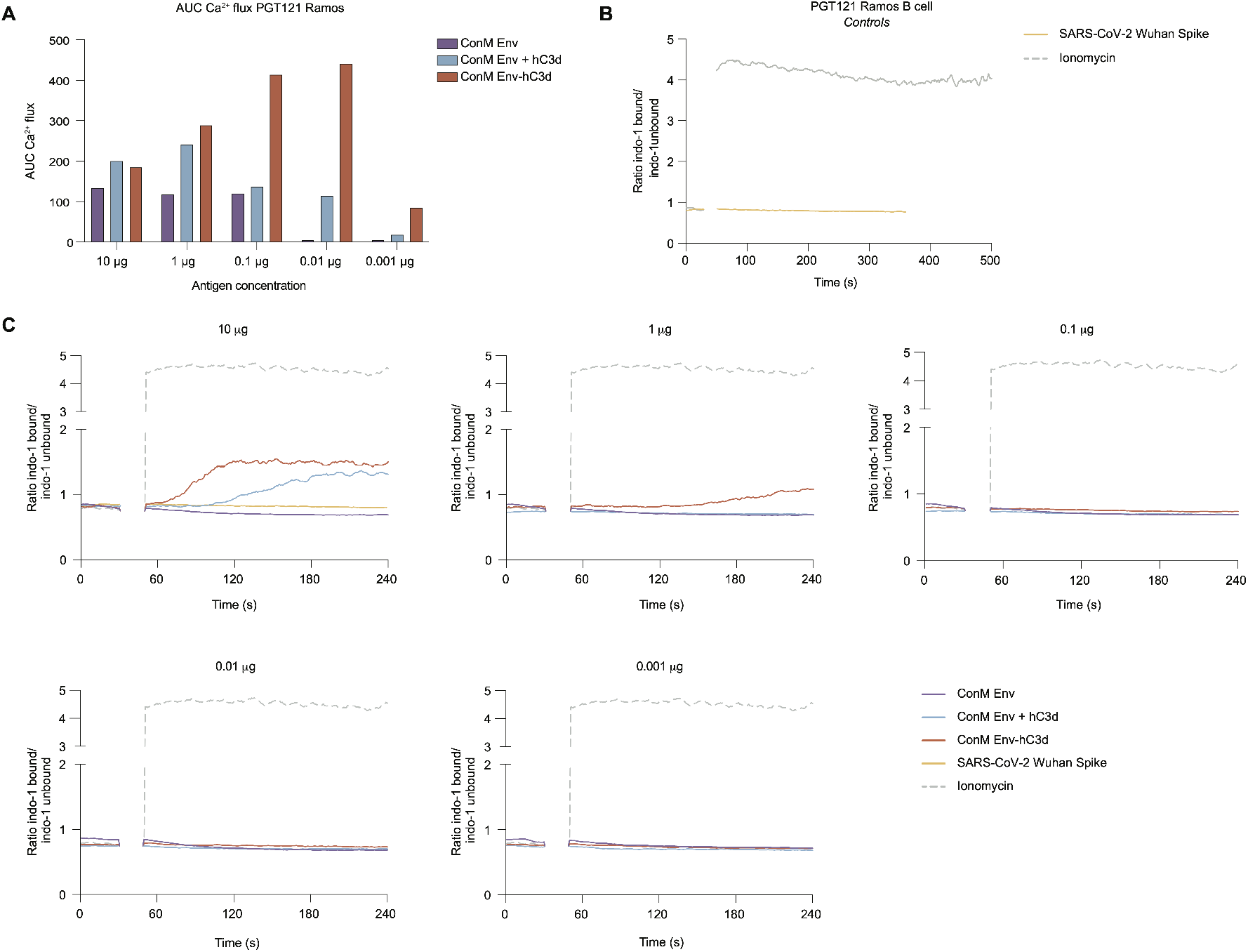
Activation of B cells expressing bNAb PGT121 or PGDM1400 by ConM Env and ConM Env-hC3d. **(A)** Area under the curve (AUC; 50–480 seconds) from calcium (Ca^2+^) flux assays shown in Figure 2. PGT121-expressing Ramos B cells were stimulated with ConM Env, ConM Env-hC3d, or ConM Env supplemented with soluble hC3d (ConM Env + hC3d) at indicated concentrations. **(B)** Ca^2+^ flux assay with PGT121-expressing Ramos B cells stimulated with 1 µg/mL ionomycin serving as a positive control to establish maximal response and 10 µg/mL of SARS-CoV-2 Wuhan Spike protein as a negative control. **(C)** Activation of PGDM1400-expressing Ramos B cell assessed through a Ca^2+^ flux assay using equimolar amounts of ConM Env, ConM Env-hC3d, or ConM Env supplemented with recombinant hC3d (ConM Env + hC3d) at concentrations of 10 µg, 1 µg, 0.1 µg, 0.01 µg, and 0.001 µg (Env-equivalent mass). ConM Env and recombinant hC3d were co-administered at a molar ratio matching the stoichiometry of ConM Env-hC3d. Ionomycin was used at 1 µg/mL as positive control and 10 µg/mL of SARS-CoV-2 Wuhan Spike as negative control. A baseline without antigen was established between 0 and 30 s, after which the measurement was interrupted to add the antigen to the B cells (30–50 s).

**Figure S3:**
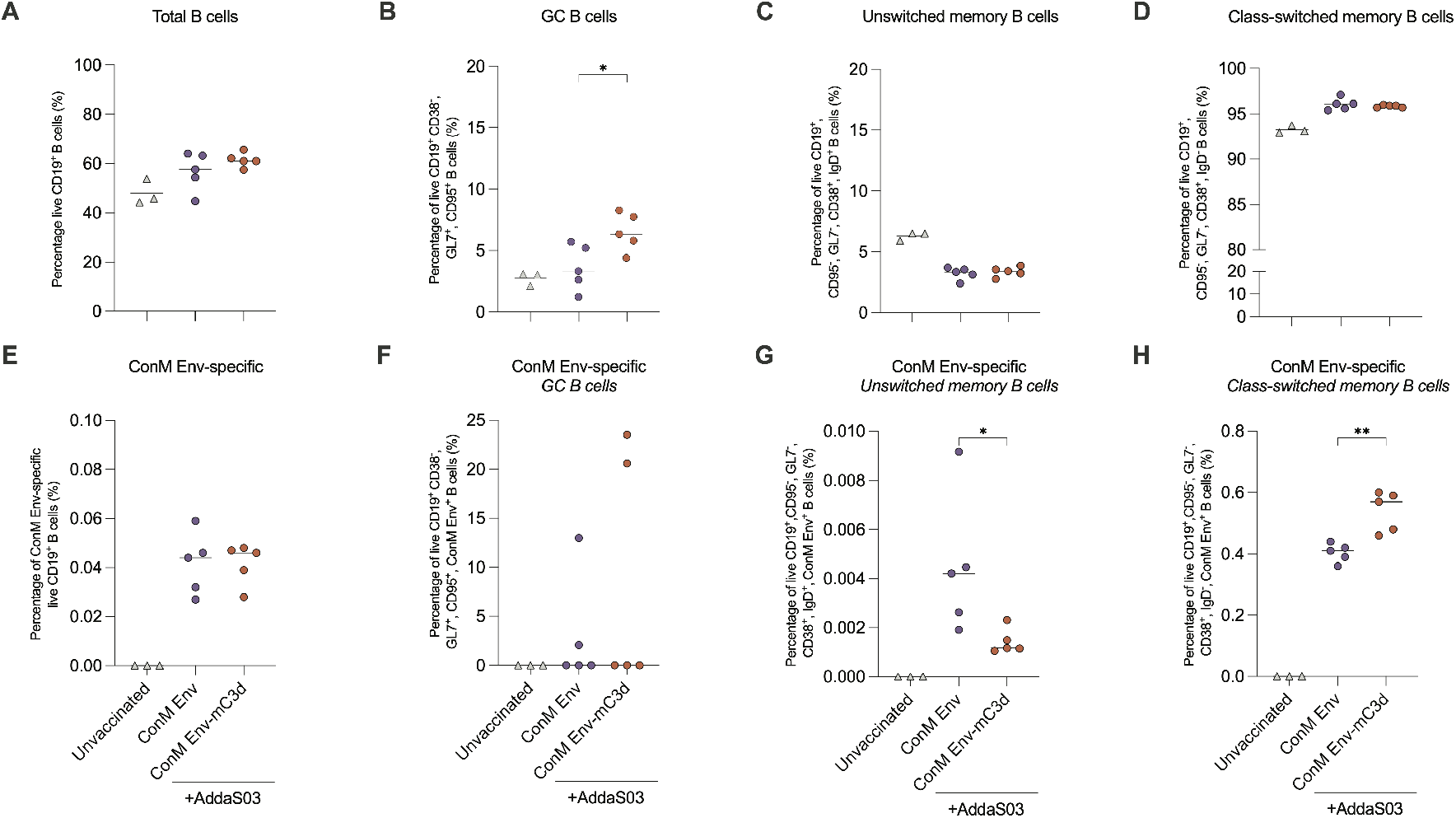
Splenic B cell populations following ConM Env or ConM Env-mC3d immunization. Mice were immunized as described in Figure 4A. Male mice (n = 5 per group) received three doses of either ConM Env or ConM Env-mC3d, each adjuvanted with AddaS03. An unvaccinated male control group (n = 3) was included for comparison. Splenocytes were harvested 4 weeks after the final immunization and analyzed by flow cytometry. **(A)** Frequency of total CD19^+^ B cells. **(B)** Frequency of ConM Env-specific B cells. **(C)** Frequency of GC B cells (CD19^+^ CD38^-^ GL7^+^ CD95^+^). **(D)** Frequency of ConM Env-specific GC B cells. **(E)** Frequency of unswitched memory B cells (CD19^+^ CD38^+^ GL7^-^ CD95^-^ IgD^+^). **(F)** Frequency of ConM Env-specific unswitched memory B cells. **(G)** Frequency of class-switched memory B cells (CD19^+^ CD38^+^ GL7^-^ CD95^-^ IgD^-^). **(H)** Frequency of ConM Env-specific class-switched memory B cells. Data are shown as scatter dot plots with the median indicated by a horizontal line. Each dot represents one mouse. Statistical comparisons between groups were performed using two-tailed Mann–Whitney U tests. Significance is indicated as: ns (p > 0.05); * (p < 0.05); ** (p < 0.01); *** (p < 0.001).

**Figure S4:**
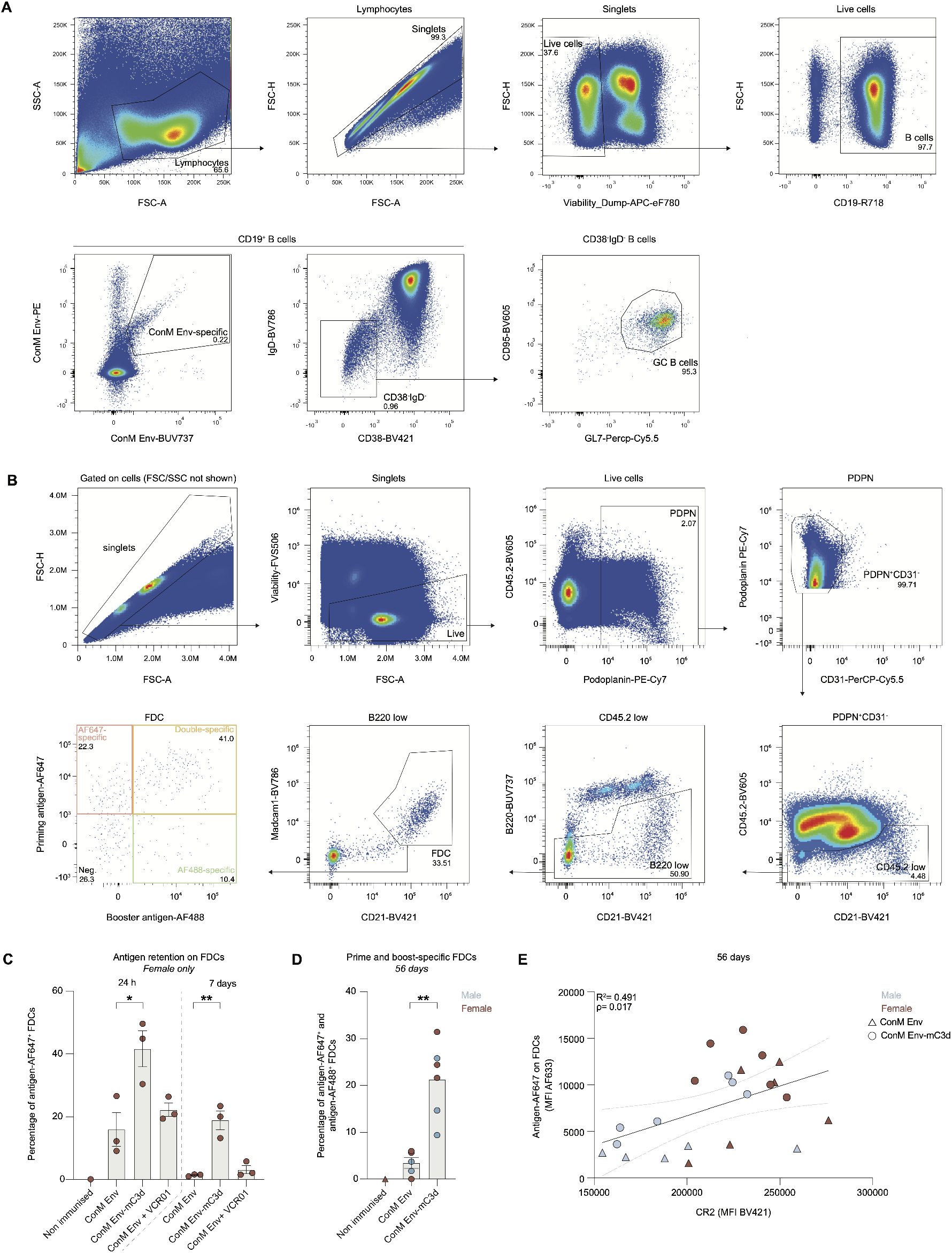
Flow cytometry gating, antigen retention, and correlation with CR2 expression in lymph node FDCs. **(A)** Gating strategy for identifying germinal center (GC) B cells and double-specific ConM Env^+^ B cells in mouse lymph nodes. First, lymphocytes were selected, followed by exclusion of doublets (FSC-H vs. FSC-A) and dead cells (viability dye). B cells were identified as CD19^+^, after which ConM Env-specific cells were gated as double-positive for BUV737- and PE-labeled ConM probes. GC B cells were defined within the CD19^+^ population as CD38^-^ IgD^-^ GL7^+^ CD95^+^. **(B)** Gating strategy for selecting FDCs from mouse lymph nodes involved excluding hematopoietic (CD45^+^ PDPN^-^) and B cells (B220^+^), while selecting stromal cells (PDPN^+^), non-endothelial (CD31^-^), that express high levels of CR2 (CD21/35^hi^) and integrin receptor Madcam1^+^. **(C)** Bar graph depicting the percentage of antigen-AF647^+^ FDCs (CD45^-^,B220^-^,CD31^-^,PDPN^+^,CD21/35^hi^,Madcam1^+^) at 24 hours and 7 days post-immunization in female mice treated with ConM Env-AF647, ConM Env-mC3d-AF647, and ConM Env-AF647 + VRC01, as described in Figure 5A. **(D)** Percentage of double-specific antigen-AF647^+^ FCDs (prime-specific) and boost antigen-AF488^+^ FDCs (boost-specific) for mice treated as described in Figure 5D, assessed at 56 days post-immunization. Sex differentiation is shown with females in maroon and males in light blue. Data are shown as bar plots with mean ± SEM. Each dot represents one mouse. Statistical comparisons were performed using two-tailed Mann–Whitney U tests. Significance is indicated as: ns (p > 0.05); * (p < 0.05); ** (p < 0.01). **(E)** Correlation between CR2 expression and antigen retention on FDCs in draining lymph nodes at 56 days post-immunization. Data from all mice were pooled, and a Pearson correlation was used to assess the relationship between CR2 expression (measured as CD21-BV421 MFI) and antigen retention on FDCs (measured as antigen-AF647 MFI); the R^2^ and p values shown reflect this analysis. Each symbol represents an individual mouse: circles indicate ConM Env-mC3d-immunized mice, and triangles indicate ConM Env-immunized mice. Blue symbols represent males, and maroon symbols represent females. The solid line indicates the best-fit linear regression; shaded dotted lines show the 95% confidence interval.

